# UltraTimTrack: a Kalman-filter-based algorithm to track muscle fascicles in ultrasound image sequences

**DOI:** 10.1101/2024.08.07.607010

**Authors:** Tim J. van der Zee, Paolo Tecchio, Daniel Hahn, Brent J. Raiteri

## Abstract

**Background:** Brightness-mode (B-mode) ultrasound is a valuable tool to non-invasively image skeletal muscle architectural changes during movement, but automatically estimating architectural features such as fascicle length remains a major challenge. Existing fascicle tracking algorithms either require time-consuming drift corrections or yield noisy estimates that require post-processing. We therefore aimed to develop an algorithm that tracks fascicles without drift and noise across a range of experimental conditions and image acquisition settings.

**Methods:** We applied a Kalman filter to combine fascicle length and fascicle angle estimates from existing and openly available UltraTrack and TimTrack algorithms into a hybrid algorithm called UltraTimTrack. We applied the hybrid algorithm to ultrasound image sequences collected from the human medial gastrocnemius of healthy individuals (*N*=8, 4 women), who performed cyclical submaximal plantar flexion contractions or remained at rest during passive ankle joint rotations at given frequencies and amplitudes whilst seated in a dynamometer chair. We quantified the algorithm’s tracking accuracy, noise, and drift as the respective mean, cycle-to-cycle, and accumulated between-contraction variability in fascicle length and fascicle angle. We expected UltraTimTrack’s estimates to be less noisy and to drift less across experimental conditions and image acquisition settings, compared with estimates from its parent algorithms.

**Results:** The proposed algorithm had low-noise estimates like UltraTrack and was drift-free like TimTrack across the broad range of conditions we tested. Estimated fascicle length and fascicle angle deviations accumulated to 2.1 ± 1.3 mm (mean ± s.d.) and 0.8 ± 0.7 deg, respectively, over 120 cyclical contractions. Average cycle-to-cycle variability was 1.4 ± 0.4 mm and 0.6 ± 0.3 deg, respectively. In comparison, UltraTrack had similar cycle-to-cycle variability (1.1 ± 0.3 mm, 0.5 ± 0.1 deg) but greater cumulative deviation (67.0 ± 59.3 mm, 9.3 ± 8.6 deg), whereas TimTrack had similar cumulative deviation (1.9 ± 2.2 mm, 0.9 ± 1.0 deg) but greater variability (3.5 ± 1.0 mm, 1.4 ± 0.5 deg). UltraTimTrack was significantly less affected by experimental conditions and image acquisition settings than its parent algorithms. It also performed well on a previously published image sequence from the human tibialis anterior, yielding a smaller root-mean-square deviation from manual tracking (fascicle length: 2.7 mm, fascicle angle: 0.7 deg) than a recently proposed hybrid algorithm (fascicle length: 4.5 mm, fascicle angle: 0.8 deg) and a machine-learning (DL_Track) algorithm (fascicle length: 8.2 mm, fascicle angle: 4.8 deg).

**Conclusion:** We developed a Kalman-filter-based method to improve fascicle tracking from B-mode ultrasound image sequences. The proposed algorithm provides low-noise, drift-free estimates of muscle architectural changes that may better inform muscle function interpretations.

## Introduction

Brightness-mode (B-mode) ultrasonography, or ultrasound, is a non-invasive method for looking under the skin to image the human body’s tissues, which can be applied to study skeletal muscle function during movement [1,2]. During passive movements and active muscle contraction, ultrasound can be used to visualize the connective tissue around muscle fascicles - bundles of muscle fibers - and their changes in length and orientation. These ‘features’ and their rates of change represent the fascicles’ behaviour which affects a muscle’s force potential [3,4] and metabolic energy expenditure [5,6], and thus can affect movement performance and economy [7,8]. Recent years have seen an increase in algorithms to automate muscle ultrasound image analysis and fascicle tracking, which had previously been limited by the laborious and subjective nature of manual labelling [9]. However, existing fascicle tracking algorithms are still error prone and have limitations that prevent complete automatization [10]. It would thus be helpful to develop an algorithm that automatically and robustly tracks muscle architectural changes from ultrasound image sequences to more quickly and easily interpret muscle function during movement.

Optical flow is a commonly used method to track muscle fascicles in a ‘semi-automated’ manner [11]. Optical-flow-based algorithms such as UltraTrack [12] make a best guess (i.e. least-squares approximation) of the apparent motion between consecutive ultrasound images to track muscle fascicles [12,13]. This method is relatively insensitive to the speckle noise present in ultrasound images because comparing two consecutive frames effectively removes common noise. However, integrating small tracking errors, which can occur because of image brightness changes or variable local pixel motion, causes fascicle length and fascicle angle estimates to ‘drift’ away from their original values over many frames. To correct for this drift, additional information (e.g. timing of external forces) is required. Additionally, if there is substantial motion between consecutive frames, then manual tracking is required to avoid underestimation of the fascicle length and fascicle angle changes. Fascicles also need to be defined before tracking, which is typically done via manual labelling. Consequently, the noise insensitivity of optical-flow-based methods is an important advantage for tracking small fascicle displacements (e.g. those that occur during healthy postural sway) [14] over a few frames, but the sensitivity of these methods to drift and manual fascicle labelling requirements reduces their accuracy, objectivity, repeatability, and time effectiveness in other conditions (e.g. large fascicle displacements during locomotion).

Unlike optical-flow-based methods, line-detection-based and artificial-intelligence (AI)-based methods attempt to automatically identify line segments in muscle ultrasound images. Line-detection-based algorithms such as TimTrack [15] analyse each ultrasound image independently, making them insensitive to drift [15–22]. However, because similarities and differences between consecutive images are not considered, these methods are sensitive to the speckle noise present in ultrasound images. This also seems to be the case for recently proposed AI-based algorithms such as DL_Track [10], which employ machine-learning techniques instead of traditional line-detection methods [10,23]. Both types of algorithms thus yield relatively noisy estimates of muscle architectural changes during movement that require filtering or averaging over multiple trials. Consequently, the automatic detection of line segments from these methods is an important advantage to increase objectivity, repeatability, and time effectiveness, but the sensitivity of these methods to noise reduces their accuracy.

To leverage the key advantages and overcome the main limitations of existing fascicle tracking algorithms, new or improved approaches are needed. Instead of developing a new method, it may be possible to combine existing algorithms using sensor-fusion techniques. Kalman filtering is a popular sensor-fusion method [24] that has been successfully employed to correct drift [e.g., 25] and reduce noise [e.g., 26] in robotics and other fields. To test whether Kalman filtering can be applied to fascicle tracking, we developed a Kalman-filter-based method. The method combines estimates from a noise-insensitive, but drift-sensitive, optical flow method (i.e. Kanade-Lucas-Tomasi optical flow) [27] with drift-free, but noise-sensitive, line-detection methods (i.e. object detection and Hough-transform methods) [28] to yield improved estimates of muscle architectural changes (Fig. 1). We re-used and modified open-source code from existing UltraTrack [12] and TimTrack [15] algorithms, and therefore coined the proposed algorithm ‘UltraTimTrack’. The proposed algorithm and its parent algorithms were tested on various B-mode ultrasound image sequences collected from the left-sided medial gastrocnemius muscle of healthy human participants. The proposed algorithm was also applied to a previously collected ultrasound image sequence from the human tibialis anterior muscle and compared with a recently proposed algorithm that combines optical-flow-based and line-detection-based algorithms, but lacks the Kalman filter (here referred to as HybridTrack) [24], as well as the AI-based DL_Track algorithm [10]. We expected UltaTimTrack to be noise-insensitive like optical-flow-based algorithms, and to be drift-free like line-detection-based and AI-based algorithms.

**Figure 1.**
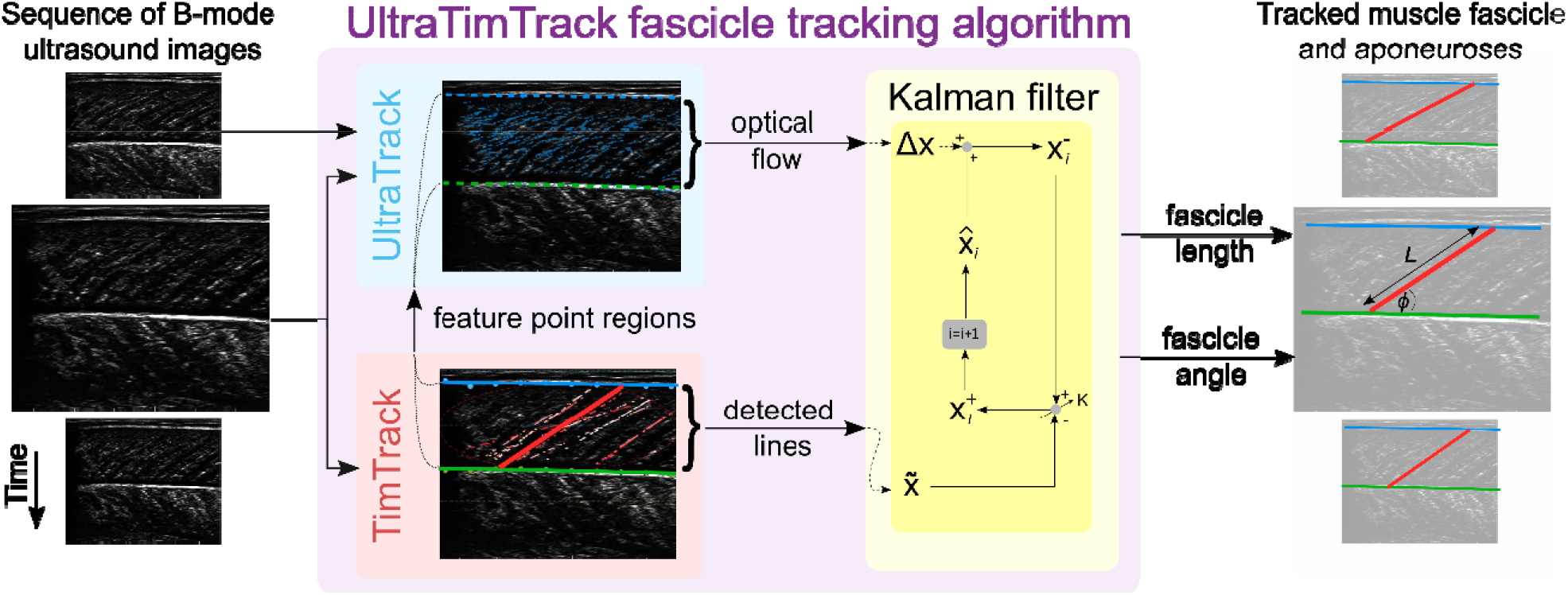
Overview of the UltraTimTrack algorithm. Left panel: Sequence of Brightness-mode (B-mode) ultrasound images. Middle panel: UltraTimTrack algorithm consists of an UltraTrack module, a TimTrack module, and a Kalman filter. First, the TimTrack module detects lines in the image corresponding to either aponeuroses or fascicles. The UltraTrack module then defines feature points near TimTrack’s detected lines and calculates the optical flow that captures the displacement of these points between the current and previous image. Optical flow and detected lines are input to the Kalman filter. In each i-th iteration, the change in estimated state is obtained from optical flow to yield an a-priori estimate, which is updated with detected lines in proportion to a Kalman gain K to yield an a-posteriori estimate. State estimates yield estimates of the tracked fascicle and its length and angle. Right panel: Tracked fascicle and aponeuroses displayed on the original images.

## Methods

### Synopsis

We developed a Kalman-filter-based fascicle tracking algorithm that combines tracking estimates from existing and openly-available algorithms to yield improved estimates of muscle fascicle length and fascicle angle changes during movement. The proposed algorithm was evaluated using ultrasound image sequences collected from the left-sided medial gastrocnemius muscle of healthy young adults during cyclical submaximal voluntary fixed-end plantar flexion contractions at various frequencies with varying activation levels, as well as during passive ankle rotations at various angular velocities (see Appendix for experimental methods). We first describe the algorithm before discussing the experimental methods, expected outcomes and statistics.

### Proposed algorithm

To improve the accuracy of the proposed algorithm, we first modified one of the parent algorithms as discussed immediately below. We then describe the Kalman filter, followed by the graphical user interface.

### Parent algorithms of UltraTimTrack

The UltraTrack and TimTrack modules of our proposed algorithm (Fig. 1) are based on the original algorithms. However, notable modifications were made to UltraTrack to improve its fascicle tracking performance. Firstly, the UltraTrack module employs Kanade–Lucas–Tomasi optical flow [30], while the original UltraTrack algorithm employs Lucas–Kanade optical flow [27]. The main difference is that Kanade–Lucas–Tomasi only tracks ‘good feature points’ that can be tracked well (in our case, corner points), which improves tracking accuracy while reducing computational cost. The Kanade-Lucas-Tomasi method has been used in more recent optical-flow-based fascicle tracking algorithms [e.g., 13], including later and improved, but non-peer-reviewed, versions of UltraTrack [31–33]. Similar to these versions, MATLAB’s (MathWorks, Inc., Natick, MA) built-in detectMinEigenFeatures, PointTracker and estimateGeometricTransform2D functions were used to detect feature points and compute optical flow with an affine transformation type, respectively. Optical flow parameters were set to values used in the most recent version of UltraTrack [31]. However, unlike this and previous UltraTrack versions, the UltraTrack module of UltraTimTrack separately computes optical flow for fascicles and aponeuroses. It leverages TimTrack’s detection of fascicle and aponeurosis locations to more precisely identify feature points that are specific to these structures. With the region of interest (ROI) type (Fig. 2) set to Hough - local (default), feature points are selected in distinct, local fascicle regions based on TimTrack’s detected fascicle locations. Specifically, the fascicle regions of the UltraTimTrack module are rectangles of fixed width (default: 10 pixels) centered around the locations of the most frequently occurring lines (default: 10) detected by TimTrack. Alternatively, ROI type can be set to Hough - global to allow feature points to be detected within the whole ROI between aponeuroses. In both cases, regions outside the middle portion (default: 80%) of the overall region between TimTrack’s aponeuroses were excluded to avoid detecting feature points on aponeuroses, similar to an existing algorithm [13]. A fixed number of feature points was selected within the remaining fascicle region (default: 300) using MATLAB’s built-in selectStrongest function. The aponeurosis regions were defined as TimTrack’s superficial and deep regions, which were specified by the user (default: 5-30% and 30-70% of vertical image range, with 0% referring to the top of the cropped ultrasound image).

**Figure 2:**
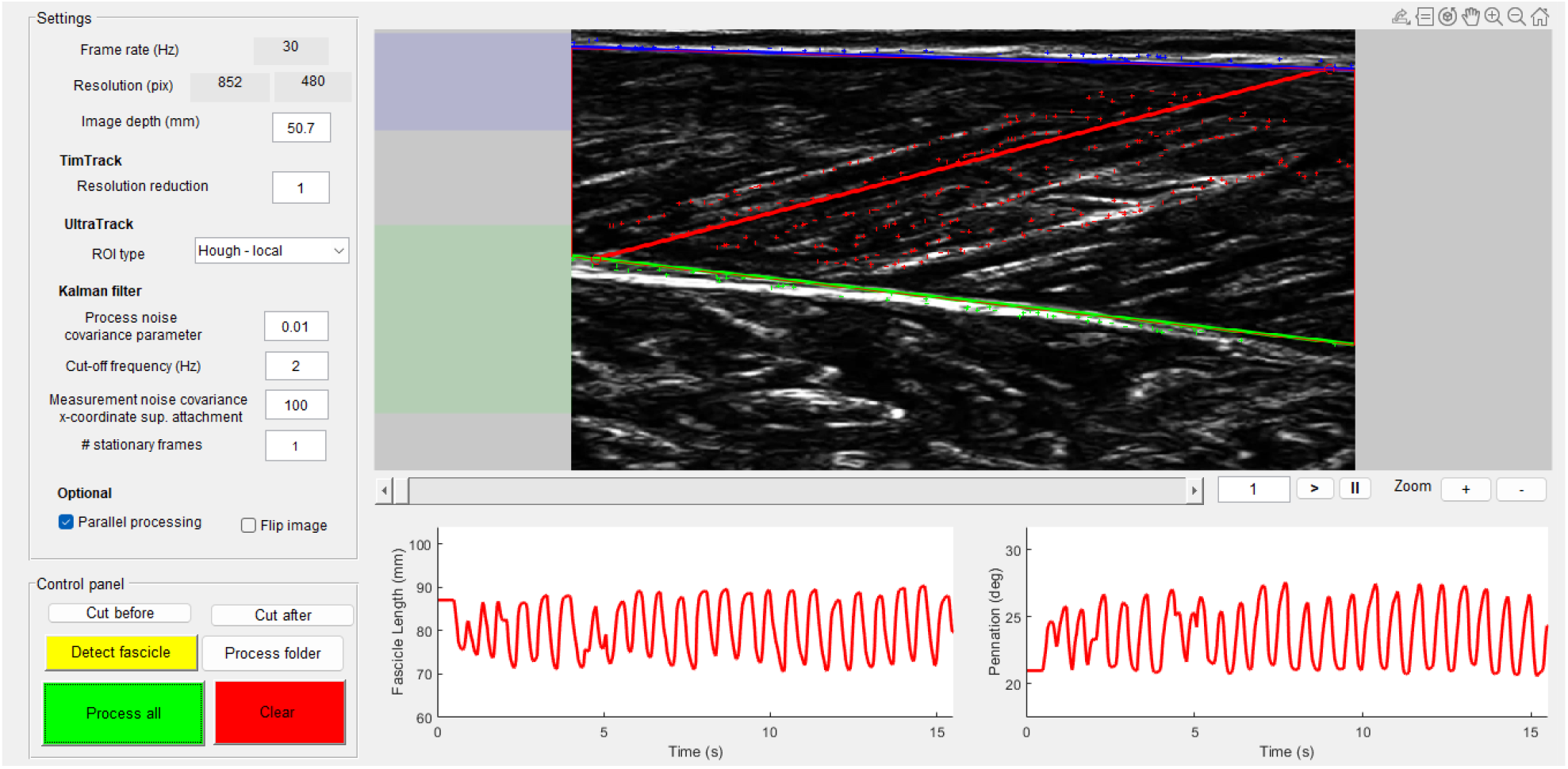
Graphical user interface of the UltraTimTrack algorithm. Users can inspect the tracked fascicle (red line) in the current ultrasound image, along with the fascicle length and fascicle angle estimates from the entire sequence. The analyzed image indicates the tracked fascicle (red line), superficial aponeurosis (blue line), deep aponeurosis (green line), and their respective feature points (red plus symbols). The aponeuroses are detected within specified regions (shaded blue and green areas to the left of the image). Users can specify the settings (top left column) and run the algorithm (bottom left column) for either the entire image sequence (Process all) or just the current frame (Detect fascicle). Settings are explained in the main text.

### Kalman filter

We now briefly describe the basics of the Kalman filter and then discuss how it was applied to track muscle fascicles. The reader is referred to other literature for a more detailed description of Kalman filters [24].

A Kalman filter assumes that the state (vector) of a system *z*_*i*_ depends on the previous state (vector) *z*_*i*−1_and input *u*:

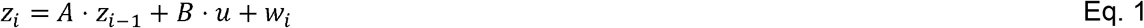

Here, *A* is a matrix that specifies the (known) state-transition model, *B* specifies the (known) control-input model and *w*_*i*_ is the (unknown) process noise, assumed to be drawn from a zero-mean normal distribution with variance *Q*_*i*_ Because the process noise *w*_*i*_ is unknown, state *z*_*i*_ cannot be computed directly. Instead, the Kalman filter can predict the state, and iteratively update this prediction to yield an optimal state estimate. To do this, the state is first predicted from a previous state estimate 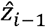 to yield an *a-priori estimate z*_*i*_ ^−^:

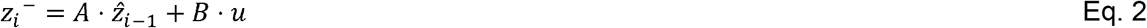

To improve the estimate, the *a-priori* state estimate *z*_*i*_ ^−^ is updated using a measurement 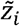:

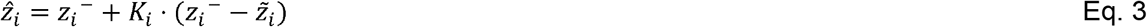

Here, *k*_*i*_ is the Kalman gain, which can be computed from the estimated noise covariances (explained later). 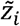 is the measurement that relates to the state *z*_*i*_:

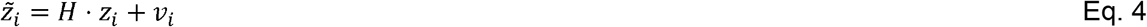

Here, *H* is the measurement model and *v* is the measurement noise, assumed to be drawn from a zero-mean normal distribution with variance *R*_*i*_. Apart from estimating the state, the Kalman filter also predicts its covariance *P*_*i*_ from the previous covariance estimate 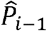 and the process noise variance *Q*_*i*_:

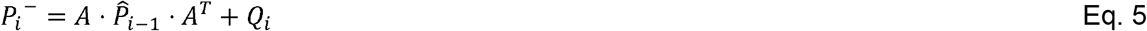

Here, *P*_*i*_ ^−^ is the *a priori* estimate of the state covariance. The Kalman gain *K*_*i*_ is computed as:

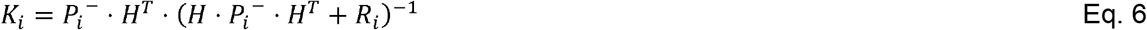

Once the Kalman gain is known, the *a-posteriori* state estimate 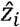 can be computed (Eq. 3), as well as its covariance 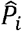:

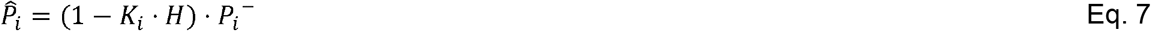

Next, we discuss the states and measurements of the UltraTimTrack fascicle tracking algorithm, and the procedure to estimate the noise covariances *Q*_*i*_ and *R*_*i*_.

### States and measurements

The proposed fascicle tracking algorithm performs Kalman filtering on both aponeuroses and fascicles. For aponeuroses, the states are the vertical locations of the superficial and deep aponeuroses at the left and right image boundaries. For fascicles, the states are the fascicle orientation and *α* the horizontal position of the fascicle’s most superficial point *p*. In both cases,

UltraTrack’s optical flow is treated as input *u*, and TimTrack’s line detection is treated as the measurement 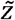. Aponeurosis locations and the fascicle orientation are updated with TimTrack’s aponeurosis locations and fascicle orientation estimate, respectively. The superficial point estimate is updated using a constant that is equal to that defined in the first frame.

### Noise estimation

To determine the Kalman gain *K* (Eq. 6), the noise covariances need to be estimated. The measurement noise covariance *R* reflects the uncertainty in TimTrack’s line detection, which is affected by image noise. We estimated the covariance of this noise by taking the variance of the high-pass filtered TimTrack aponeurosis locations and fascicle orientations. A second-order dual-pass Butterworth filter was used (default cut-off frequency: 2 Hz). The measurement noise covariance of the horizontal position of the fascicle’s superficial point reflects how well the initial position value reflects the instantaneous position value. A large covariance allows more movement of this point, but at the cost of more drift (default: 1000 pixels). Considering that optical flow is most accurate for small displacements, the process noise *Q* was assumed to increase with the square of the optical flow input *u*:

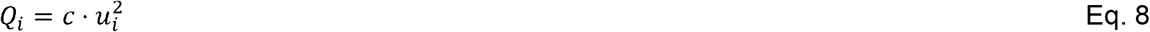

Note that unlike the measurement noise *R* that is assumed to be constant over an image sequence, the estimated process noise *Q* is different for each image *i* because the optical flow *u*_*i*_ is different. The proportionality constant *c* is unknown, and it is chosen through trial-and-error.

### Backwards smoothing: Rauch-Tung-Striebel

After Kalman filtering, smoothing may be applied to the state estimates 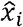 and covariance 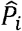 using a Rauch-Tung-Striebel smoother:

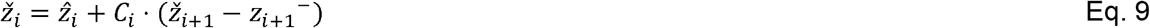

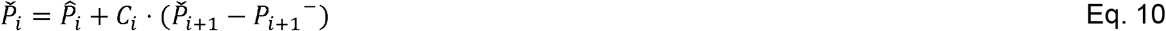

Here, 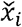 and 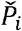 are the smoothed state estimate and covariance, respectively. The smoothing gain *C*_*i*_ is computed from the *a-priori* and *a-posteriori* covariances:

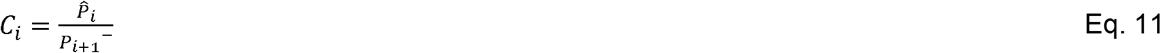

### Initial state estimate

The initial state estimate 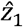 is derived from TimTrack. Optionally, the user can specify a number of frames over which the mean TimTrack value is taken from the first frame to this number for the initial estimate (default: 1), and the variance is taken as an estimate of the initial covariance 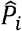. These frames must be stationary (e.g. at a steady-state passive or active muscle force during no muscle length change). If no stationary frames are available, the initial state estimate 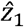 is determined for the first frame and the covariance is assumed to equal the measurement noise *R*.

### Fascicle tracking estimates

Two main fascicle tracking estimates are derived from our proposed method: (1) fascicle angle *ϕ* and (2) fascicle length *L*. These estimates are determined by 11 variables in the algorithm, of which six are states and five are constants (see Table 1).

**Table 1.** Variables of the UltraTimTrack algorithm.

| **Symbol** | **Type** | **Meaning** |
| --- | --- | --- |
| $S_{L,x}$ | Constant | Horizontal coordinate of the superficial aponeurosis left image boundary |
| $S_{L,y}$ | State | Vertical coordinate of the superficial aponeurosis left image boundary |
| $S_{R,x}$ | Constant | Horizontal coordinate of the superficial aponeurosis right image boundary |
| $S_{R,y}$ | State | Vertical coordinate of the superficial aponeurosis right image boundary |
| $D_{L,x}$ | Constant | Horizontal coordinate of the deep aponeurosis left image boundary |
| $D_{L,y}$ | State | Vertical coordinate of the deep aponeurosis left image boundary |
| $D_{R,x}$ | Constant | Horizontal coordinate of the deep aponeurosis right image boundary |
| $D_{R,y}$ | State | Vertical coordinate of the deep aponeurosis right image boundary |
| $p_x$ | State | Horizontal coordinate of the superficial fascicle point |
| $p_y$ | Constant | Vertical coordinate of the superficial fascicle point |
| $\alpha$ | State | Fascicle orientation (i.e., with respect to the horizontal) |

First, the horizontal (*x*) and vertical (*y*) coordinates of the fascicle attachment points on the superficial and deep aponeuroses ([*s*_*x*_,*s*_*y*_] and [*d*_*x*_,*d*_*y*_], respectively) are determined from the states. First, the vertical positions of the superficial aponeurosis, deep aponeurosis, and fascicles are expressed as functions of the horizontal position *x*, yielding *S*(*x*),*D*(*x*) and *F*(*x*), respectively:

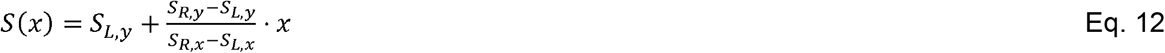

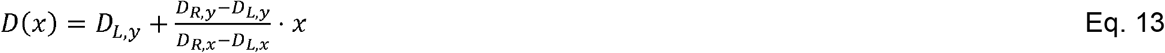

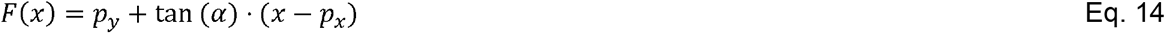

Next, the superficial and deep attachment points [*s*_*x*,_*s*_*y*_] and [*d*_*x*,_*d*_*y*_] are found by solving *F*(*x*)=*S* (*x*) and *F*(*x*)=*D*(*x*), respectively. The attachment points and aponeurosis locations yield estimates of fascicle angle *ϕ*(i.e., with respect to the deep aponeurosis) and length *L*:

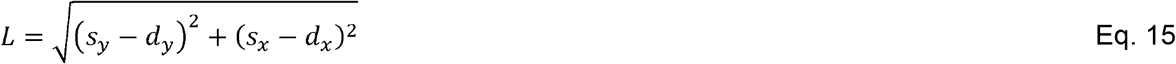

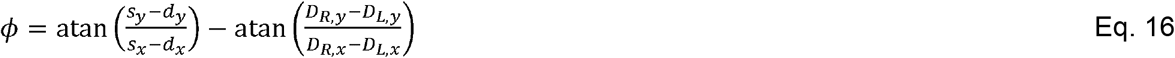

For UltraTrack, the second term of Eq. 16 was set to 0 because UltraTrack requires additional inputs to track aponeuroses.

### Graphical user interface

UltraTimTrack has a similar graphical user interface (GUI) as UltraTrack (see Fig. 2). It displays the current ultrasound image and the tracked muscle fascicle, alongside the estimated fascicle length and fascicle angle for the image sequence. The algorithm can be started using the “Process all” button and reset using the “Clear” button. The “Detect fascicle” detects a single fascicle using TimTrack for the current image, and the “Process folder” button allows all videos in a specified folder to be processed. Settings include the frame rate and resolution (obtained from the video file) and the image depth to scale pixels to mm (note that if a TVD file is converted to an MP4 file using the provided function, TVD2ALL.m, this information is stored in a MAT file and automatically loaded along with the video). The image resolution can be optionally decreased with a specified factor (“Resolution reduction”) to speed up TimTrack. Under Processing, the user can specify the estimated optical flow process noise covariance parameter (see “Noise estimation”), the cut-off frequency of the high-pass filtering, the superficial attachment point measurement noise covariance, and the number of stationary frames (see “Initial state estimate”). Under “Options”, the user can also check whether to perform parallel processing to speed up computation, and whether to flip the image about its middle vertical axis. Currently, only forward tracking from the first frame of the image sequence is supported.

### Experimental methods

To test the UltraTimTrack algorithm, ultrasound images from the left-sided medial gastrocnemius muscle of healthy human participants were collected during various active and passive conditions performed within a dynamometer setup.

#### Participants

We collected data from healthy young adults (N = 8, four women, age = 26 ± 3 years, height = 174 ± 6 cm, weight = 70 ± 7 kg) during ankle plantar flexion contractions of various speeds, various hold durations, and submaximal voluntary intensities. Participants were included based on the following criteria: age 18 – 45 years, no pre-existing neuromuscular conditions, and no injuries of the lower extremity in the last six months. The latter was determined by a standardised questionnaire that were approved by the Local Ethics Committee. Participants provided written informed consent prior to their voluntary participation in the experiment. The experiment was approved by the Ethics Committee of the Faculty of Sport Science at Ruhr-University Bochum (reference: EKS V 19/2022).

#### Experimental setup

Participants were seated on a dynamometer (IsoMed2000, D&R Ferstl GmbH, Germany) chair, with the sole of their left foot flush against the footplate attachment of a motorized dynamometer while their left knee was extended. Their left foot had an attachment above it that was padded to limit forefoot movement. A figure of eight strap around the footplate attachment and participant’s lower shank was used to avoid heel lift. The back of the dynamometer chair was reclined to an angle of about 60 deg to avoid stretch of the hamstring muscles.

Net ankle joint torque and crank-arm angle were measured using the dynamometer and sampled at 2 kHz using a 16-bit analog-to-digital converter (Power1401-3) and Spike2 (10.10 64-bit version) data collection system (Cambridge Electronic Design Ltd., Cambridge, United Kingdom), which had a ± 5 V input range.

The left-sided medial gastrocnemius muscle was imaged using a linear, flat, 128-element ultrasound transducer (LV8-5N60-A2, Telemed) that was attached to a PC-based ultrasound beamformer (ArtUS EXT-1H, Telemed). Ultrasound images were captured with a 60 × 50 mm (width x depth) field of view at frame rates ranging from 33 images·s^-1^ to 100 images·s^-1^, and image collection was synchronized with the collection of torque and crank-arm angle as previously described [31,33].

The activity of the following four lower leg muscles was recorded using surface electromyography (EMG): medial gastrocnemius, lateral gastrocnemius, soleus, and tibialis anterior. EMG was recorded using a NeuroLog system (NL905, Digitimer Ltd., Welwyn Garden City, United Kingdom). The signals were recorded at 2 kHz and synchronised using the previously described analog-to-digital converter and software.

#### Experimental conditions

The experiment involved three distinct types of conditions: (1) ∼3-s duration maximal voluntary fixed-end contractions (MVCs) of plantar flexion and dorsiflexion at 0 deg plantar flexion (sole of foot perpendicular to shank); (2) ten prolonged (80 s) bouts of submaximal voluntary fixed-end plantar flexion contractions at different rates at 0 deg plantar flexion, and; (3) passive ankle joint rotations over a 50 deg range of ankle angles at different velocities.

During the MVCs, the participants were verbally encouraged by the experimenter to push or pull with the ball or top of their foot as much as possible for at least 3 s. After each MVC, participants rested for at least two minutes. For the plantar flexion MVCs, this procedure was repeated until the maximal torque of the subsequent contractions differed by less than 5%, or until five contractions were performed, whichever came first. Following the plantar flexion MVCs, participants performed at least one dorsiflexion MVC.

During the prolonged submaximal voluntary contractions, the participant was asked to match their net ankle joint plantar flexion torque to a visually displayed torque target for a duration of 80 s in six sub-conditions. In four of those conditions, the target was to ramp up and down from 0-50% MVC and there was a 1-sec hold phase at 50% MVC. Three of the four conditions differed in that the ramp rates varied (20% MVC·s^-1^, 33% MVC·s^-1^ and 100% MVC·s^-1^) and each condition was repeated once to allow for two ultrasound image qualities (so-called “high” and “low” quality images, which were obtained with line density software settings of high and standard S, respectively) to be assessed, which resulted in at least six prolonged contraction bouts. To reduce the chance of fatigue, participants were given at least two minutes of rest between bouts. The fourth ramp-and-hold condition involved a different ramp rate up (20% MVC s^-1^) and ramp rate down (the instruction was to relax as fast as possible), and this asymmetric ramp was repeated to allow two ultrasound image qualities to be assessed, which resulted in two prolonged contraction bouts. In the remaining two conditions, the target trace was a sinusoid with a frequency of 1.5 Hz that was not repeated (only a high-image quality was used), which resulted in two prolonged contraction bouts. The two sinusoidal conditions differed in their torque range (0-20% MVC and 10-20% MVC).

Following the cyclical contractions, the participant was asked to sit still and relax while the experimenter triggered the dynamometer to passively rotate their foot eight times over 50 deg within the participant’s ankle joint range of motion. Three passive rotation conditions were performed, each at a different angular velocity: 5 deg·s^-1^, 30 deg·s^-1^, and 120 deg·s^-1^. The corresponding maximum and minimum ankle angles were 41.6 ± 7.8 deg, 41.6 ± 7.9 deg and 41.7 ± 7.8 deg plantar flexion, and 5.9 ± 6.8 deg, 5.9 ± 6.7 deg and 6.0 ± 6.8 deg dorsiflexion (mean ± s.d. across participants).

#### Experimental procedure

Before performing the experiment, participants were prepared for the EMG and ultrasound measurements. First, we located the muscly belly borders of the left-sided medial gastrocnemius using ultrasound. To improve sound wave transmission, ultrasound gel was applied over the participants’ skin. The desired location (i.e. generally the most prominent bulge of the muscle) of the ultrasound transducer over the muscle belly was marked using a red permanent marker. Next, the EMG electrodes were placed over the skin covering the four muscles of interest according to SENIAM guidelines [34]. For the medial gastrocnemius, the location was shifted medially or laterally in case it overlapped with the desired location of the ultrasound transducer. For each of the four muscles, an area of about 3-4 cm^2^ was shaved, exfoliated and disinfected before electrode placement. EMG electrodes were placed with an interelectrode center-to-center distance of 2 cm following SENIAM recommendations [34]. Next, the ultrasound transducer was secured to the lower leg using a custom-made case and self-adhesive bandage. Next, the dynamometer chair was set up, and the dynamometer axis of rotation was aligned to the participant’s ankle joint axis of rotation (estimated as the transmalleolar axis) during a 50% MVC contraction at 0 deg plantar flexion. Movement of the dynamometer footplate was restricted to occur within the participant’s ankle joint range of motion using both electrical and mechanical stops. After set-up was complete, participants were asked to perform the three distinct experimental conditions described above.

#### Data processing

Second-order dual-pass Butterworth filters were used to low-pass filter the torque and crank-arm angle data (cut-off: 10 Hz), and to band-pass filter the EMG data (cut-off: 30-500 Hz) [35]. EMG data were then rectified and low-pass filtered (cut-off: 10 Hz), again using a second-order dual-pass Butterworth filter. The filtered EMG data were normalized with respect to the corresponding maximum value obtained during the MVC trials. Finally, filtered torque, angle and EMG data were resampled to a frequency of 100 Hz.

## Expected outcomes

### Comparison with UltraTrack and TimTrack

UltraTimTrack was compared to UltraTrack and TimTrack algorithms as available from open-access GitHub repositories (i.e. github.com/brentrat/UltraTrack_v5_3 and github.com/timvanderzee/ultrasound-automated-algorithm, respectively). Combining the experimental conditions allowed us to investigate the effect of three types of factors on fascicle tracking estimates: (1) sequence duration; (2) image-to-image dissimilarity, and; (3) image quality. UltraTrack was expected to be sensitive to sequence duration, as drift accumulates over time. Ultratrack was also expected to be sensitive to image-to-image dissimilarity, because optical flow estimates are more accurate when consecutive images are more similar. TimTrack was expected to be primarily affected by image quality, because its sensitivity to noise implies that it may perform better on high-quality images. UltraTimTrack’s Kalman-filter-based method was expected to exploit UltraTrack’s and TimTrack’s complementary dependencies. Specifically, UltraTimTrack was expected to be less sensitive to sequence duration and to image-to-image dissimilarity than UltraTrack, and less sensitive to image quality than TimTrack. We used default values for all algorithm parameters, except for a resolution reduction of 2.

### Outcome measures

We estimated tracking accuracy, noisiness, and drift using a combination of objective measures and comparison to manual labelling (i.e. manual tracking).

#### Overall variability – Sinusoids and ramp-and-hold contractions

Estimated fascicle lengths and fascicle angles (i.e. features) from each contraction cycle were resampled as a percentage of contraction cycle (1% spacing). Overall variability was estimated as the mean standard deviation of the estimated fascicle features across contraction cycles:

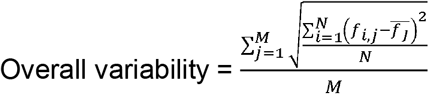

Here, *f* is the fascicle feature estimate (either fascicle length or fascicle angle), *N* is the number of contraction cycles, and *M* is the number of percentage points (set to 101, for 1% spacing). A portion of this variability reflects ‘true’ variability (e.g., due to torque-matching inaccuracies during cyclical contractions), with the remainder due to tracking errors of the algorithm. The difference in variability provides an estimate of algorithm error when comparing algorithms on the same ultrasound image sequences. This overall variability measure should increase with both drift and noisiness of each algorithm’s estimates.

#### Overall variability – Passive rotations

We used a different variability measure for the passive ankle rotations because the cycle time and number of cycles were variable. We binned each fascicle feature estimate into 1-degree joint angle bins and computed the standard deviation for each bin. Overall variability was defined as the average of these standard deviations among joint angle bins.

#### Cycle-to-cycle variability - Sinusoids

The difference in estimates between two consecutive contraction cycles was determined for each resampled point and for each pair of consecutive cycles, yielding a two-dimensional difference matrix *δ* (*M*-by-[ -1]). A mean difference was computed for each cycle pair, which yielded a mean vector 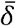(1-by-[ *N*-1]). Cycle-to-cycle variability of each algorithm’s tracking estimates was quantified using the standard deviation of the cycle-to-cycle difference in fascicle feature estimates, and averaged over contraction cycles:

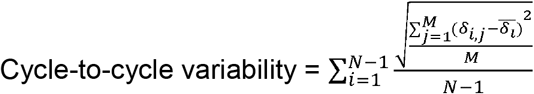

Because this measure only compares two consecutive contraction cycles, it does not capture the drift that happens over longer time scales. Instead, it mostly reflects noisiness and is less affected by drift.

#### Cumulative deviation - Sinusoids

Cumulative deviation of the tracking estimates was quantified using the cumulative rectified mean cycle-to-cycle difference of each algorithm’s fascicle tracking estimates. Cumulative deviation was evaluated at the final contraction cycle:

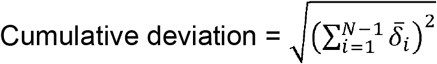

As this measure only compares the mean difference between contraction cycles, it does not reflect noisiness over short time scales (i.e. within a cycle). Instead, it mostly reflects drift and is less affected by noisiness.

### Comparison to HybridTrack and DL_Track

We also evaluated the UltraTimTrack algorithm on an example image sequence from the human tibialis anterior that accompanied the HybridTrack algorithm publication [29]. In addition to tracking this video with HybridTrack, we also tracked it with the recently proposed DL_Track AI-based algorithm [10]. We compared tracking estimates from all three algorithms to estimates from manual observers (N = 3) for 100 equidistantly-spaced images in the sequence (i.e. every 6^th^ image). Custom MATLAB code was used to manually track selected images in sequential order. Manual observers had several years of training and experience in ultrasonography and manual tracking. Processing time was also compared between algorithms, and a sensitivity analysis was performed on the effect of the unknown process noise covariance parameter *c*.

We used default values for all algorithm parameters, except for a process noise covariance parameter *c* of 0.001.

### Statistics

We evaluated fascicle tracking noise and drift for algorithm estimates from the larger-range sinusoidal trial (i.e. 0-20% MVC) using cycle-to-cycle variability and cumulative deviation measures, respectively. These measures were statistically compared between algorithms using paired t-tests with Holm-Bonferroni corrections for comparing multiple algorithms. The effect of sequence duration was tested by comparing overall variability from the 10^th^ contraction cycle to overall variability from the final contraction of the sinusoidal trials. The effect of image-to-image dissimilarity was tested by comparing the change in overall variability with (1) a larger torque range during sinusoidal trials, (2) faster ramp rates during ramp-and-hold trials, and (3) faster rotation rates during passive trials. The effect of image quality was tested by comparing overall variability between low- and high-image quality sequences of the ramp-and-hold trials. Linear mixed-effects regression models (MATLAB’s *fitlme*) were used to test these effects, using participants as a random effects variable to account for between-participant differences. Results are provided as mean ± standard deviation (s.d.), with s.d. referring to between-participants variability unless stated otherwise.

## Results

Participants produced an average ankle plantar flexion MVC torque of 114.5 ± 47.7 N·m at 0 deg plantar flexion, and produced mean torques similar to the mean target torques (Table A1) in the sinusoidal trials and ramp-and-hold trials (Fig. 3). Muscle activity changed cyclically over time (Fig. A1), similarly to torque (Table A1). As expected during the sinusoidal trials, UltraTrack’s estimates drifted, TimTrack’s fascicle estimates were noisy, and UltraTimTrack’s estimates were insensitive to both drift and noise (Fig. 3A). Similar results were obtained from the ramp-and-hold (Fig. 3B) and passive trials (Fig. 4).

**Figure 3:**
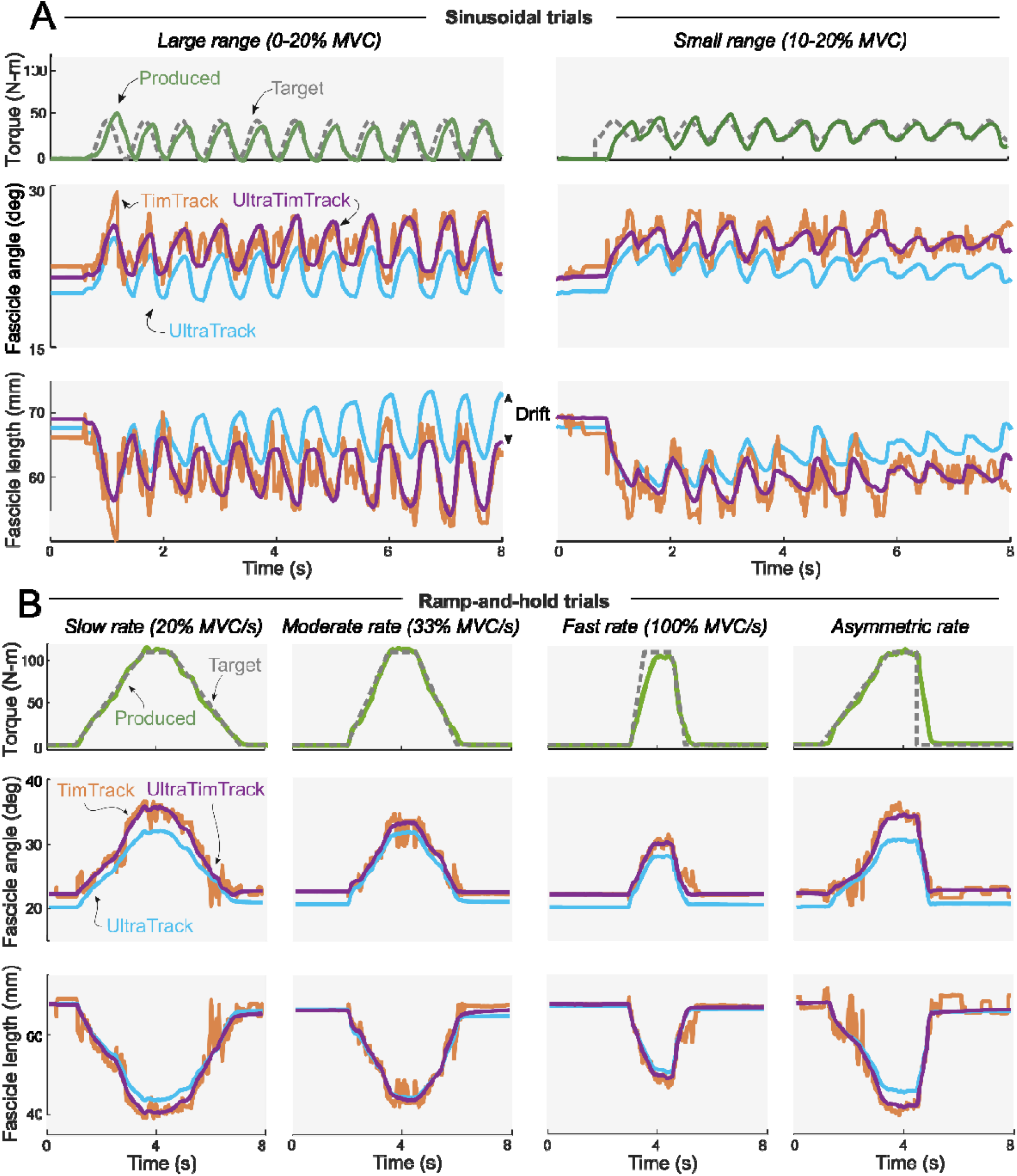
Typical example of active plantar flexion torques and fascicle tracking outputs during sinusoidal trials and ramp-and-hold trials. Participants produced cyclical torques at given frequencies and ranges for a duration of 80 s (first 8 s shown). Produced torque (green solid line) resembled the desired torque (grey dotted line). TimTrack (red), UltraTrack (blue) and the proposed UltraTimTrack (purple) algorithms yielded estimates of fascicle length and fascicle angle. **A**. Sinusoidal trials with a large torque range (left) and a small torque range (right) were performed. UltraTrack’s drift was most apparent for the fascicle length output during the large torque range trial (indicated with a double-sided arrow). TimTrack’s noisiness for both fascicle tracking outputs was apparent throughout. UltraTimTrack fascicle tracking outputs were low-noise and drift-free. **B**. Ramp-and-hold trials with different ramp rates (slow, moderate, fast, asymmetric) were performed. UltraTrack’s fascicle angle estimates appear offset because it does not consider the angle of the aponeurosis; TimTrack’s estimates were noisy. MVC: Maximal Voluntary Contraction.

**Figure 4:**
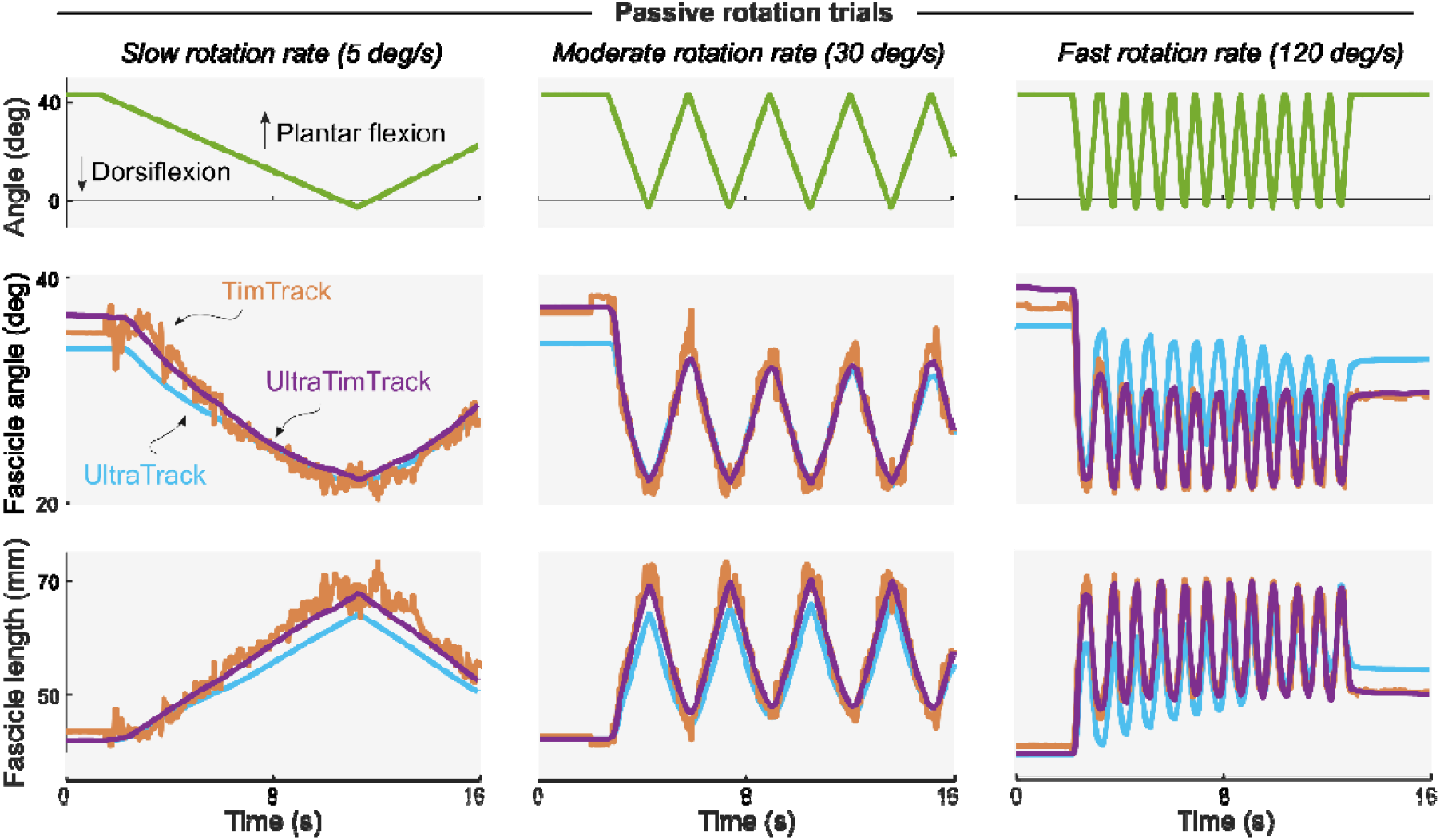
Typical example of passive plantar flexion torques and fascicle tracking outputs during passive rotation trials. Columns show trials with different passive rotation rates (slow, moderate, and fast). Participants relaxed while the experimenter rotated their left ankle joint through its range of motion (first 16 s shown). TimTrack (red), UltraTrack (blue) and the proposed UltraTimTrack (purple) algorithms yielded estimates of fascicle length and fascicle angle. UltraTrack’s fascicle angle estimates were offset because it does not consider the angle of the aponeurosis; TimTrack’s estimates were noisy.

UltraTimTrack’s estimates had lower cycle-to-cycle variability and cumulative deviation than its parent algorithms (Fig. 5). Cycle-to-cycle variability of fascicle length and fascicle angle estimates of UltraTimTrack (1.4 ± 0.4 mm and 0.6 ± 0.3 deg) was smaller than TimTrack (3.5 ± 1.0 mm and 1.4 ± 0.5 deg; p = 0.001 and p = 0.002, respectively, paired t-tests with Holm-Bonferroni corrections), but not different from UltraTrack (1.1 ± 0.3 mm and 0.5 ± 0.1 deg; p = 0.082 and p = 0.195, respectively). UltraTimTrack’s cumulative deviation of fascicle length and angle estimates during sinusoidal contractions between 0-20% MVC (2.1 ± 1.3 mm and 0.8 ± 0.7 deg) was smaller (p = 0.018 and p = 0.028, respectively) than UltraTrack’s (67.0 ± 59.3 mm and 9.3 ± 8.6 deg), but not different from TimTrack (1.9 ± 2.2 mm and 0.9 ± 1.0 deg, p = 0.623 and p = 0.476, respectively). Unlike its parent algorithms, UltraTimTrack had both low cycle-to-cycle variability and low cumulative deviation, indicating insensitivity to noise and drift, respectively.

**Figure 5:**
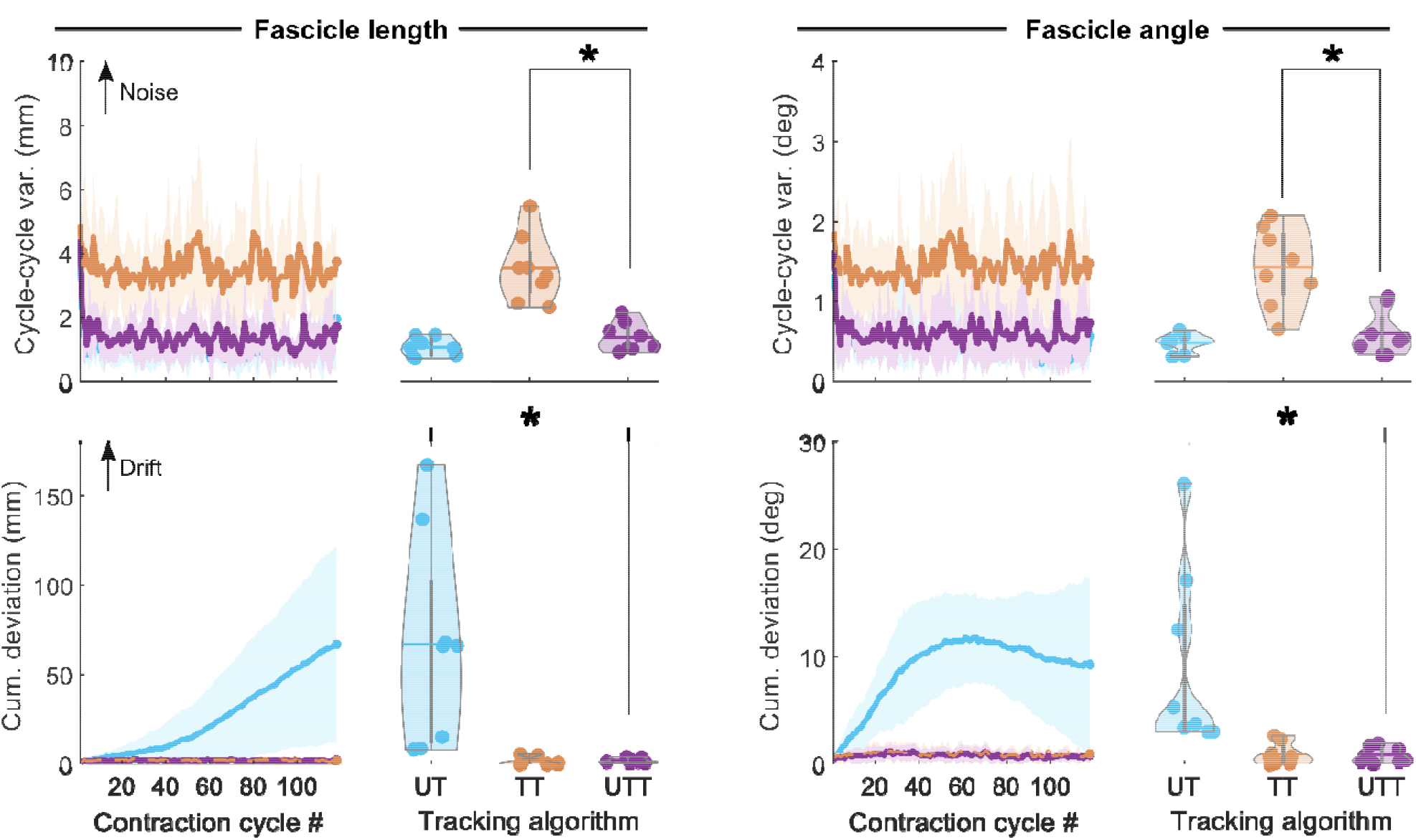
Noise and drift of the proposed and parent fascicle tracking algorithms. Data are shown for sinusoidal contractions with a large torque range (0-20% MVC), and for fascicle length (left) and fascicle angle (right). Top row: Cycle-to-cycle variability (var.) of all contraction cycles and averaged over contraction cycles. Cycle-to-cycle variability is sensitive to noise. Bottom row: Cumulative (cum.) deviation of all contraction cycles and the last cycle. Cumulative deviation is sensitive to drift. UltraTimTrack (UTT, purple) had lower cumulative deviation than UltraTrack (UT, blue), and lower cycle-to-cycle variability than TimTrack (TT, red); asterisks indicate significant differences (p < 0.05). Lines indicate mean across participants; shaded area indicates mean ± standard deviation.

Overall variability of fascicle tracking estimates from UltraTimTrack was generally comparable to or lower than its parent algorithms, and less sensitive to sequence duration, and image-to-image dissimilarity, and image quality (Fig. 6). For sinusoidal trials, UltraTimTrack was less sensitive to both the sequence duration and image-to-image dissimilarity than UltraTrack, as indicated by significant interaction effects between the number of contraction cycles or torque range with algorithm type on fascicle length variability (algorithm [cycle number]: p = 0.004, algorithm [torque range]: p = 0.009, linear mixed-effects regression with repeated measures). Similar interactions were found for fascicle angle variability (algorithm [cycle number]: p = 9·10^−7^, algorithm [torque range]: p = 0.006). There were no significant interactions with algorithm type when comparing UltraTimTrack to TimTrack for fascicle length variability (algorithm [cycle number]: p = 0.363, algorithm [torque range]: p = 0.464) or fascicle angle variability (algorithm [cycle number]: p = 0.409, algorithm [torque range]: p = 0.105). However, UltraTimTrack had lower overall variability than TimTrack as indicated by independent effects of algorithm (fascicle length: p = 0.0003, fascicle angle: p = 0.0004). For symmetric ramp-and-hold trials, UltraTimTrack was less sensitive to image-to-image dissimilarity than UltraTrack, as indicated by interactions between ramp rate and algorithm type on estimate variability (fascicle length: p = 0.015, fascicle angle: p = 4·10^−7^). There was no significant interaction between image quality and algorithm type on estimate variability (fascicle length: p = 0.829, fascicle angle: p = 0.997), which indicates that UltraTimTrack and UltraTrack had a similar sensitivity to image quality. UltraTimTrack had lower overall variability than TimTrack as indicated by independent effects of algorithm (fascicle length: p = 1·10^−12^, fascicle angle: p = 5·10^−11^). UltraTimTrack was more sensitive to image quality for fascicle length estimates (algorithm × [image quality]: p = 0.035), but not for fascicle angle estimates (p = 0.089), and more sensitive to image-to-image dissimilarity for fascicle angle estimates (algorithm × [ramp rate]: p = 0.009), but not for fascicle length estimates (p = 0.097). For the asymmetric ramp trial, UltraTimTrack had a lower fascicle length and angle variability than both UltraTrack (p = 0.0001 and p = 0.0002) and TimTrack (p = 0.001 and p = 0.0002), with no differences in sensitivity to image quality (algorithm × [image quality]: p’s > 0.1). For the passive trials, UltraTimTrack was less sensitive to image-to-image dissimilarity than UltraTrack for fascicle length (algorithm × [rotation rate]: p = 0.044) and angle estimates (p = 0.009). There was no significant difference in sensitivity to image-to-image dissimilarity between UltraTimTrack and TimTrack for fascicle length (algorithm × [rotation rate]: p = 0.055) or angle estimates (p = 0.327), but overall variability was lower for UltraTimTrack than TimTrack (fascicle length: p = 2·10^−8^, fascicle angle: p = 5·10^−6^). Overall, UltraTimTrack had similar or lower estimate variability than its parent algorithms across a range of conditions, indicating both accurate and robust fascicle tracking.

**Figure 6:**
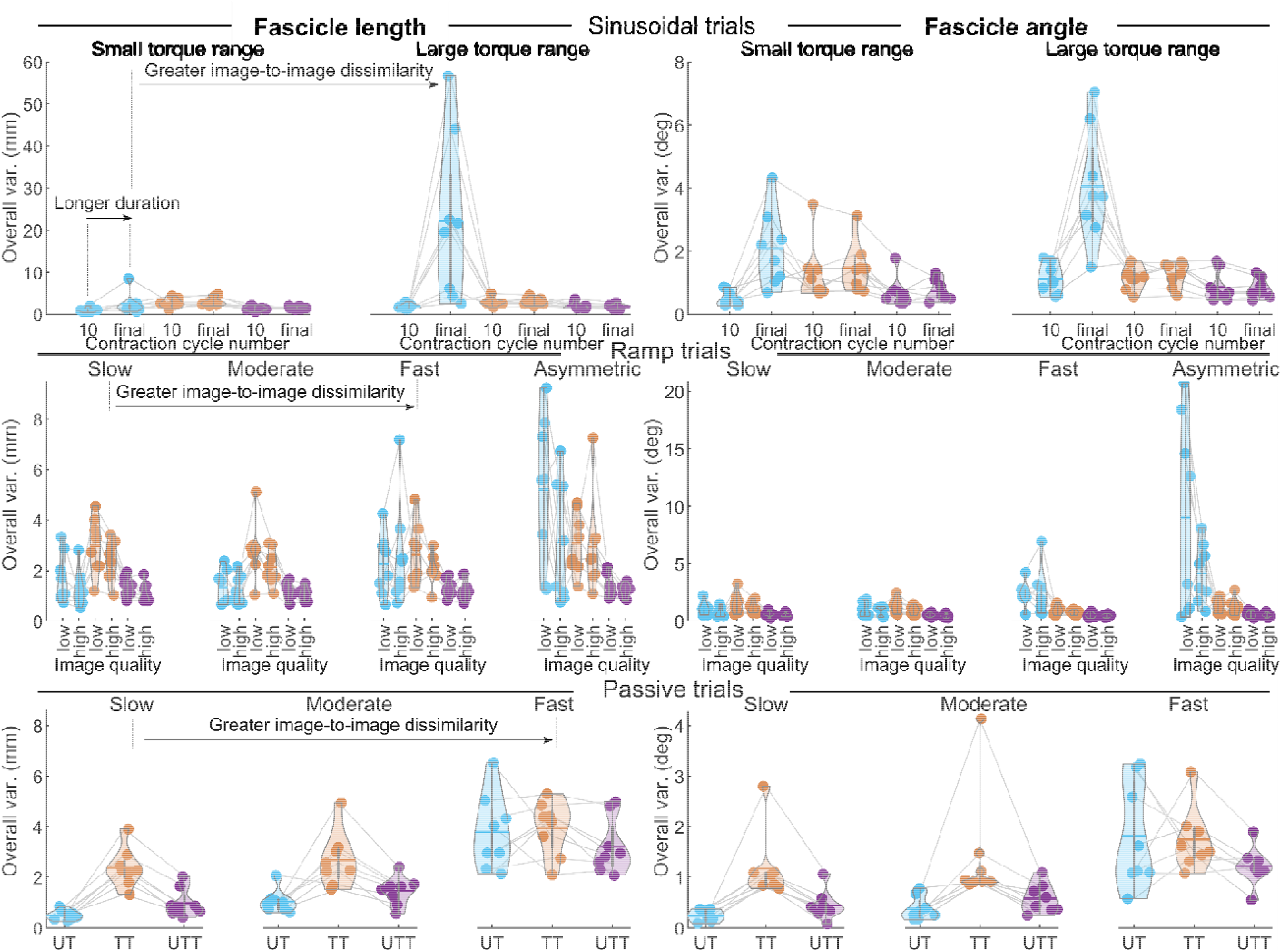
Overall estimate variability of the proposed and parent fascicle tracking algorithms. Ultrasound tracking algorithm estimates of fascicle length (left) and fascicle angle (right) are shown for all experimental conditions. Trials belonged to one of three categories: sinusoidal (top row), ramp-and-hold (middle row) or passive rotation (bottom row). Within categories, three aspects of the ultrasound image sequences varied: (1) sequence duration (sinusoidal trials only), (2) image-to-image dissimilarity (all trials), or (3) image quality (ramp-and-hold trials only). For most trials, overall variability of UltraTimTrac (UTT, purple) was similar to or lower than that of either UltraTrack (UT, blue) or TimTrack (TT, red), and similarly or less sensitive to settings during ultrasound image collection. Statistics are provided in the text.

UltraTimTrack also compared favorably against recently proposed fascicle tracking algorithms HybridTrack and DL_Track algorithms for the example video from the human tibialis anterior muscle that accompanied the HybridTrack publication [29] (see Fig. 7). Despite notable out-of-plane motion in this video, both UltraTimTrack and HybridTrack yielded low-noise (i.e. smooth) and drift-free fascicle tracking estimates, but with different amplitudes. UltraTimTrack yielded a smaller root-mean-square deviation (RMSD) relative to manual estimates (fascicle length: RMSD = 2.7 mm, fascicle angle: RMSD = 0.7 deg) compared with HybridTrack (fascicle length: RMSD = 4.5 mm, fascicle angle: RMSD = 0.8 deg) and DL_Track (fascicle length: RMSD = 8.2 mm, fascicle angle: RMSD = 4.8 deg). UltraTimTrack yielded smaller RMSD’s than HybridTrack for a range of estimated process noise covariance parameter values *c* spanning about 4 orders of magnitude (see Fig. A2). Videos of fascicle tracking by UltraTimTrack and HybridTrack are available as Supplemental Information. UltraTimTrack’s processing time (124 s, Intel Core i7-10700 CPU @ 2.90GHz) was 4.8 times smaller than HybridTrack (597 s), and 8.4 times smaller than DL_Track (1046 s). When parallel processing was employed (8 cores), processing time decreased further (72 s, 8.3 - 14.5 times faster). UltraTimTrack thus performs well with a lower computational cost compared with openly available state-of-the-art algorithms.

**Figure 7:**
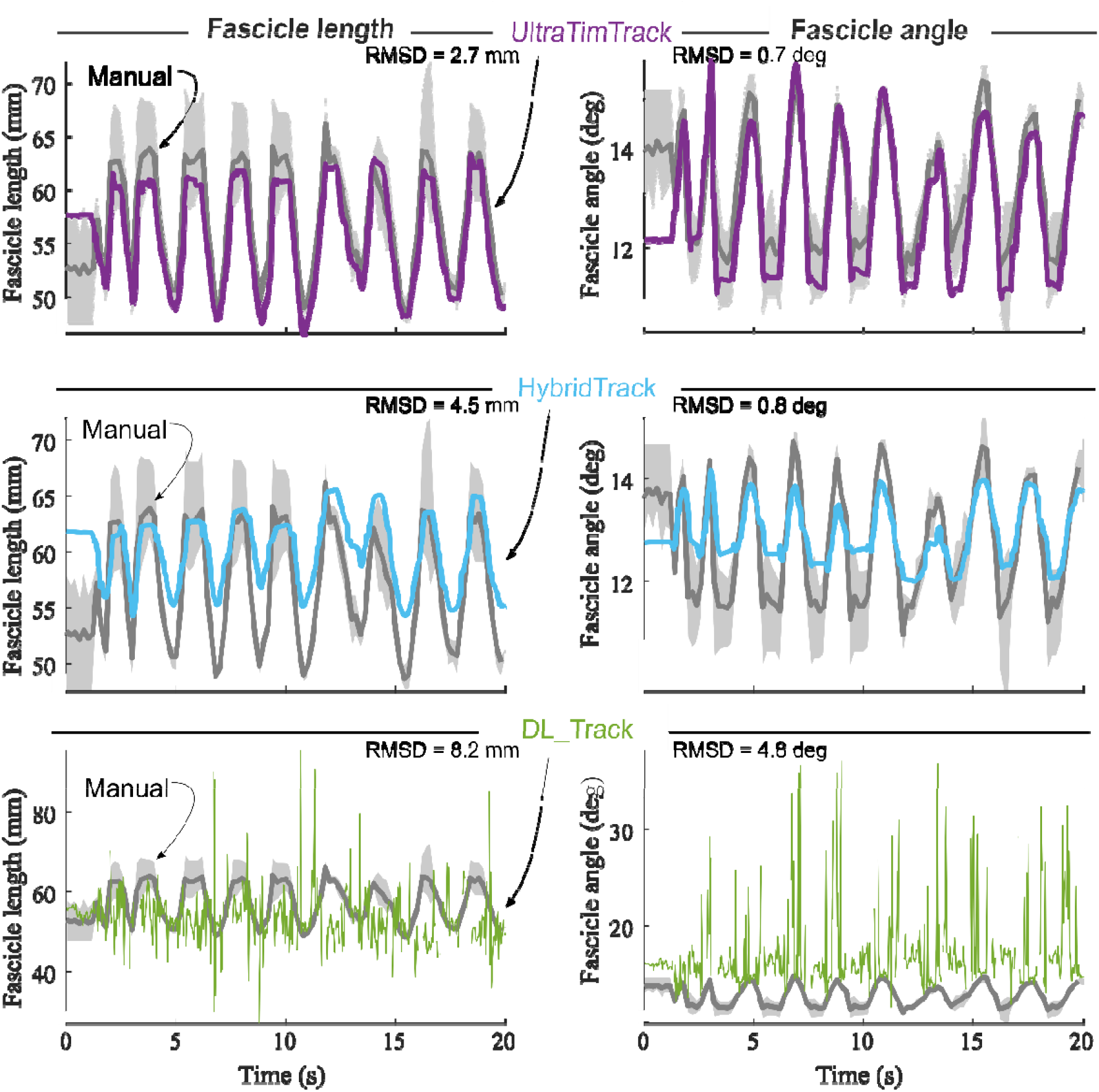
UltraTimTrack and recently proposed fascicle tracking algorithms compared with manual tracking. A published ultrasound image sequence from the human tibialis anterior muscle during fixed-end dorsiflexion contractions was tracked with the proposed UltraTimTrack algorithm (purple, top row), with HybridTrack (blue, middle row), with DL_Track (green, bottom row), and manually by three independent observers (grey, all rows). Fascicle length estimates (left column) and fascicle angle estimates (right column) were compared between algorithms with the mean outputs from manual tracking using the root-mean-square deviation (RMSD). HybridTrack estimates were obtained from the associated publication that included this image sequence. UltraTimTrack had a smaller RMSD from manual tracking estimates than either previously published tracking algorithm. Manual tracking outputs are shown as mean (thick grey lines) and range (light grey shaded areas) across observers. Note that the manual tracking outputs are the same for each algorithm comparison but appear different because of different scaling of the vertical axis to accommodate each algorithm’s outputs.

## Discussion

We proposed a Kalman-filter-based fascicle tracking algorithm that combines drifting estimates of fascicle length and fascicle angle from optical-flow methods with noisy estimates of fascicle angle from line-detection methods to improve muscle fascicle tracking. The proposed method employs existing UltraTrack and TimTrack algorithms as modules for optical flow estimation and line detection, respectively, and combines outputs from both algorithms in the UltraTimTrack algorithm to reduce subjectivity and the time burden of fascicle tracking. Unlike existing algorithms, UltraTimTrack’s fascicle length and fascicle angle outputs are low-noise and drift-free and are obtained at a low computational cost.

The proposed fascicle tracking algorithm yielded estimates without the drift of UltraTrack and without the noise of TimTrack. Less noise and drift were observed for time series from both of a representative participant (Fig. 3-4) and from participant-average summary measures (Figs. 5-6). Unlike TimTrack, UltraTimTrack exploits information from optical flow in the Kalman-filter method to reduce noise. This allows UltraTimTrack estimates to be smooth even for data from a single contraction. UltraTrack also uses this optical flow information, but without an automatic update step to correct drift. UltraTrack’s estimates therefore drift with each contraction, causing its contraction average to be offset compared with drift-free algorithms (Fig. 3). Averaged over participants, UltraTimTrack had less drift than UltraTrack and less noise than TimTrack (Fig. 5). Less noise and drift resulted in similar or low overall variability of UltraTimTrack’s estimates across contraction cycles compared with its parent algorithms (Fig. 6). Furthermore, the proposed Kalman-filter-based method was generally less sensitive to factors that are known to affect tracking performance, including sequence duration, image-to-image dissimilarity, image quality. UltraTimTrack thus provides robust estimates of fascicle length and angle changes during movement compared with existing algorithms.

The proposed fascicle tracking algorithm also performed well compared with more recently proposed and openly available fascicle tracking algorithms. Like UltraTimTrack, HybridTrack’s estimates appeared low-noise and drift-free, albeit with different amplitudes of fascicle length and angle changes (Fig. 6). We believe that HybridTrack may have a smaller amplitude because it (1) filters the Hough angle quite heavily, and (2) it uses the similarity transform instead of the affine transform in optical flow. Filtering of Hough angles included a smoothing spline curve, and a 10-point moving average. Employing a similarity transform instead of an affine transform may yield a smaller fascicle length change amplitude because shear is neglected, which can be an important contributor to fascicle-length and fascicle-angle changes [36]. We would like to point out that this may be an error in HybridTrack, because in this publication it is mentioned that an affine transformation was used. UltraTimTrack’s estimates agreed more closely with manual tracking, suggesting better accuracy. DL_Track’s estimates deviated more from manual tracking than either UltraTimTrack or HybridTrack. DL_Track’s relatively large deviations may be due to a combination of sensitivity to image noise, and an inability to generalize beyond the data it was trained on. DL_Track’s sensitivity to noise is apparent in the accompanying publication [10] and may originate from the fact that it tracks multiple (and different) fascicles in each image. Because of the heterogeneity between line segments of the same and different fascicles, this can cause a noisy estimate of the dominant fascicle angle and subsequently of fascicle length. This noise sensitivity was likely exacerbated by using ultrasound images that were different from those used to train DL_Track. DL_Track could not detect a single fascicle or the deep aponeurosis in some images and misidentified the aponeurosis in others. Its performance may be expected to improve when optimizing some built-in parameters and when re-trained on data more similar to the data considered here, but this would require (1) those data being available, and (2) manual labelling. Alternatively to manual labelling, it may be useful to input UltraTimTrack’s low-noise and drift-free fascicle tracking outputs. However, it remains to be seen whether AI-based algorithms can be trained enough to work for any ultrasound image sequence in a plug-and-play manner. Apart from a higher accuracy, UltraTimTrack’s processing time was also smaller than both HybridTrack and DL_Track, with time savings of approximately 5-6 and 6-11 times, respectively. Overall, UltraTimTrack appears to have clear advantages compared with currently available state-of-the-art fascicle tracking algorithms in terms of fascicle tracking accuracy and computational cost.

Ultrasound-based estimates of muscle architecture have many potential applications in muscle physiology and movement science fields. Musculoskeletal simulations are popular methods for studying movement and commonly use Hill-type muscle models to predict muscle excitations, forces, lengths, and even movement itself [37]. Such simulations may benefit from ultrasound-derived estimates of muscle architecture, which could be used to fit model parameters [38] or test model predictions [39]. Considering the low computational cost and high apparent accuracy of the proposed algorithm (<0.15 s per image), ultrasound-derived estimates may also be used as real-time feedback to help control movements, either by a user or an external controller. The feasibility of such a ‘muscle-in-the-loop’ approach was previously proposed for a machine-learning algorithm [23], but it has not been implemented yet for this purpose to our knowledge.

UltraTimTrack has limitations that could be addressed in future updates of the algorithm. For example, the algorithm assumes that fascicles and aponeuroses are straight lines, while they may be curved in certain cases (e.g. at rest and low activation levels). Another limitation is that both the process noise covariance parameter *c* and superficial aponeurosis measurement noise covariance parameter are unknown. It may be better to inform parameter *c* based on independent data rather than through trial-and-error. Furthermore, we assumed a constant measurement noise covariance, although it may be expected to vary between images. In future versions of the algorithm, measurement noise may be estimated for each image individually using the variance of a certain number of Hough angles.

## Conclusion

We developed a Kalman-filter-based fascicle tracking algorithm that combines existing optical-flow-based and line-detection-based algorithms to yield low-noise and drift-free fascicle length and fascicle angle estimates from B-mode ultrasound image sequences. The proposed algorithm has a low computational cost and may be adapted for real-time fascicle tracking and for training AI-based approaches. UltraTimTrack is openly available to facilitate its improvement and to allow others to utilize its benefits and make modifications to address their specific needs.

## Appendix

**Table S1:**
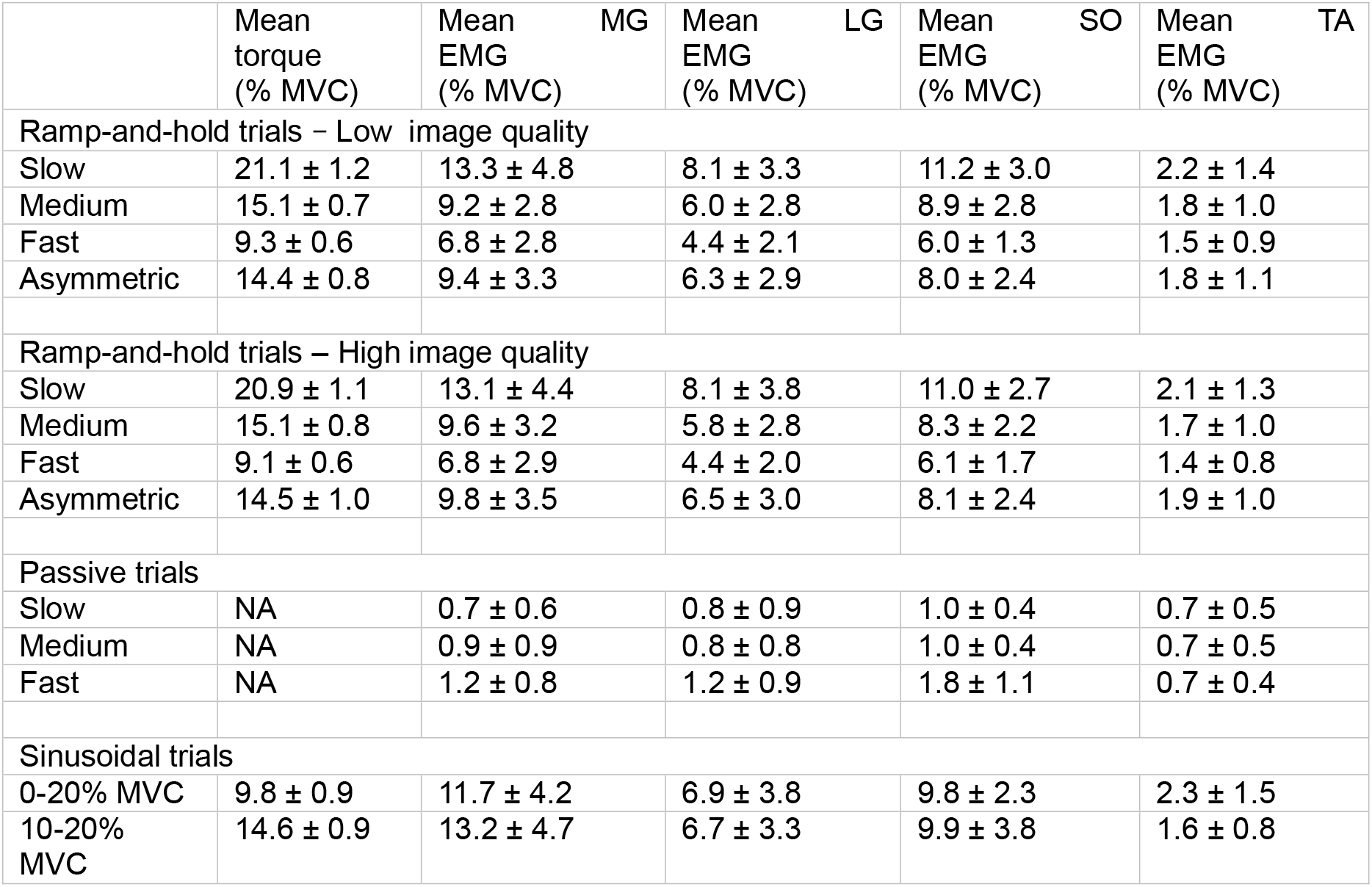
Mean torques and EMG amplitudes

**Fig. S1:**
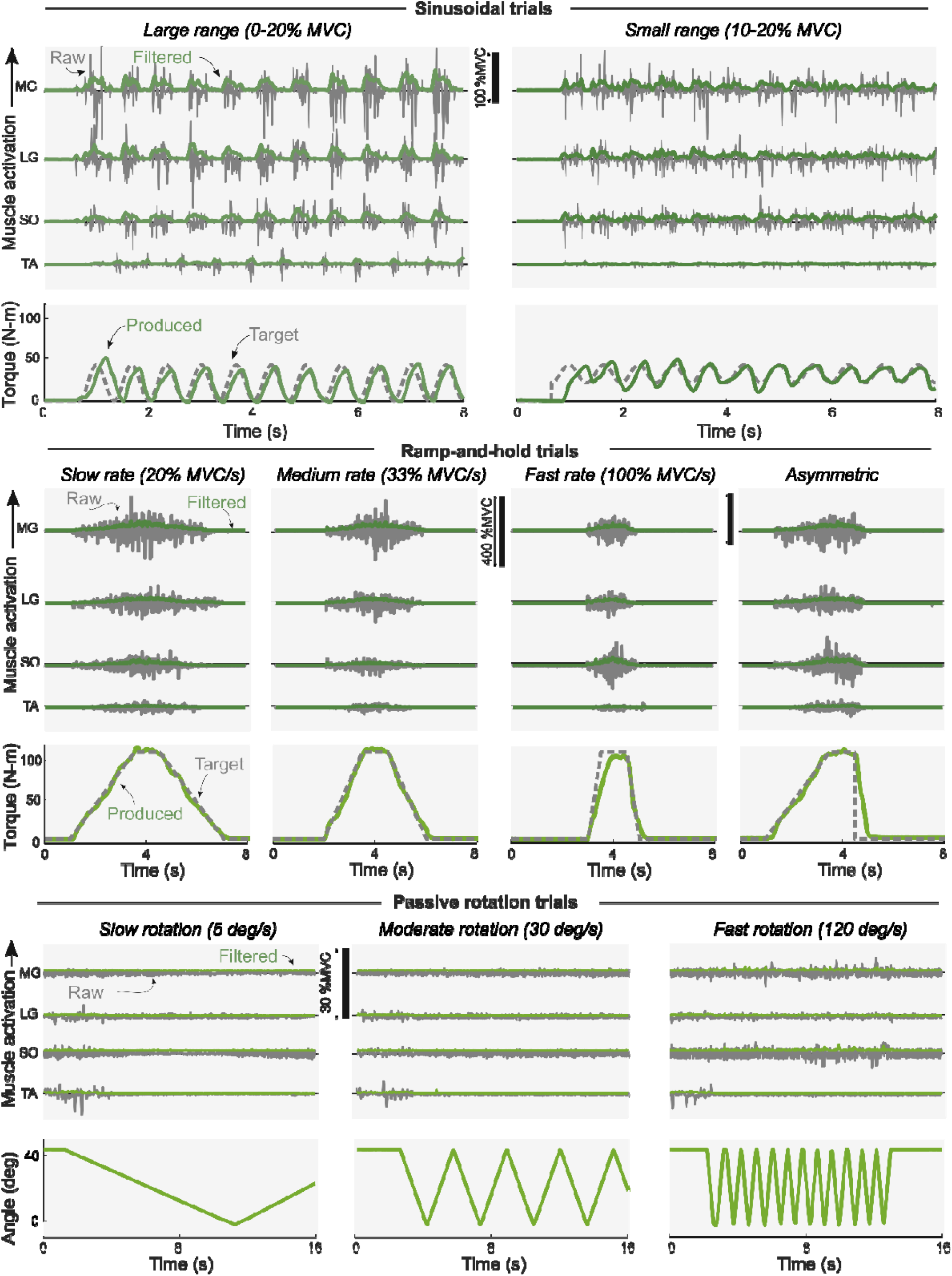
**Typical example of muscle activity levels during the three distinct experimental conditions.**

**Fig. S2:**
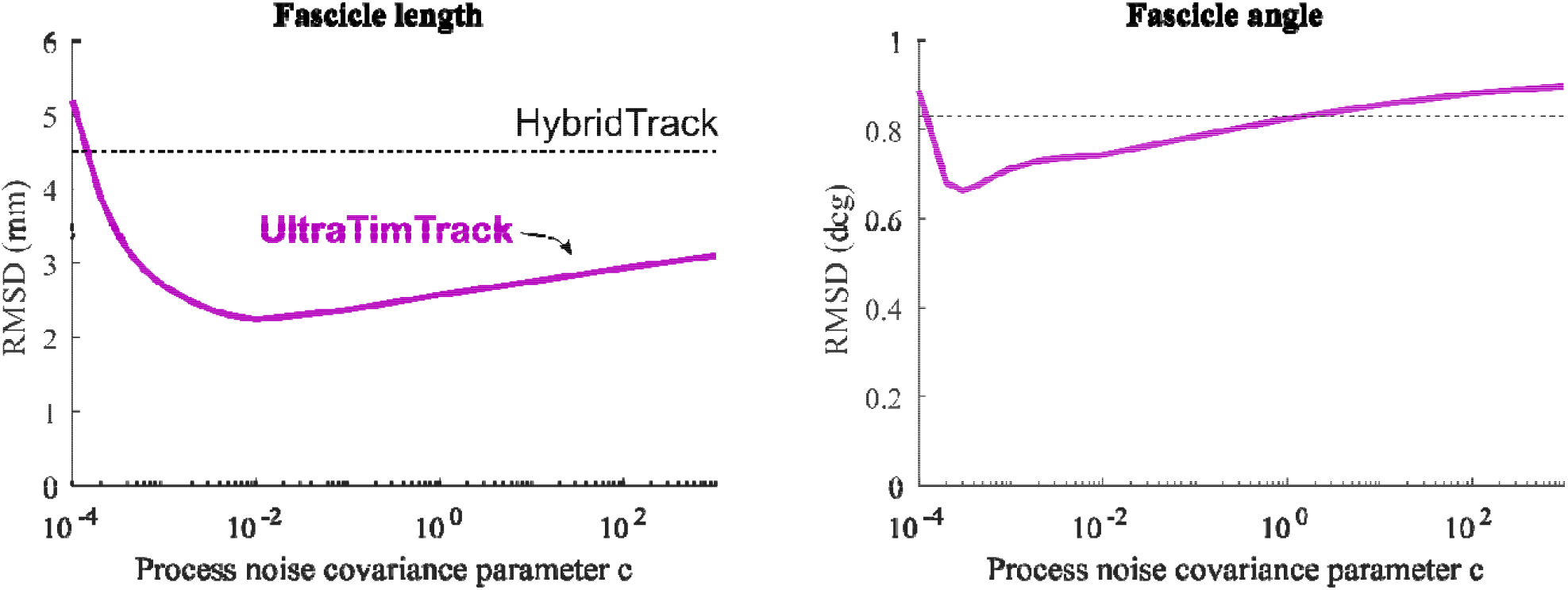
Sensitivity analysis on covariance parameter. A published ultrasound image sequence from the human tibialis anterior muscle during fixed-end dorsiflexion contractions was tracked with the proposed UltraTimTrack algorithm (purple solid line) and with HybridTrack (black dotted line). Process noise covariance parameter was varied, while all other parameter values were kept constant. Root-mean-square deviation (RMSD) of fascicle length (left) and fascicle angle (right) from manual tracking of human observers (N = 3) changed less than 3-fold for a range of values spanning 7 orders of magnitude.

## References

[1] O.M. Rutherford, D.A. Jones, Measurement of fibre pennation using ultrasound in the human quadriceps in vivo, Eur J Appl Physiol Occup Physiol 65 (1992) 433–437. 10.1007/BF00243510.

[2] T. Fukunaga, K. Kubo, Y. Kawakami, S. Fukashiro, H. Kanehisa, C.N. Maganaris, In vivo behaviour of human muscle tendon during walking, Proc. R. Soc. Lond. B 268 (2001) 229–233. 10.1098/rspb.2000.1361.

[3] S. Bohm, F. Mersmann, A. Santuz, A. Arampatzis, The force–length–velocity potential of the human soleus muscle is related to the energetic cost of running, Proceedings of the Royal Society B: Biological Sciences 286 (2019) 20192560. 10.1098/rspb.2019.2560.

[4] E. Azizi, E.L. Brainerd, T.J. Roberts, Variable gearing in pennate muscles, PNAS 105 (2008) 1745–1750. 10.1073/pnas.0709212105.

[5] O.N. Beck, L.H. Trejo, J.N. Schroeder, J.R. Franz, G.S. Sawicki, Shorter muscle fascicle operating lengths increase the metabolic cost of cyclic force production, Journal of Applied Physiology (2022). 10.1152/japplphysiol.00720.2021.

[6] V. Joumaa, W. Herzog, Energy cost of force production is reduced after active stretch in skinned muscle fibres, Journal of Biomechanics 46 (2013) 1135–1139. 10.1016/j.jbiomech.2013.01.008.

[7] W. Swinnen, I. Mylle, W. Hoogkamer, F. DE Groote, B. Vanwanseele, Changing Stride Frequency Alters Average Joint Power and Power Distributions during Ground Contact and Leg Swing in Running, Med Sci Sports Exerc 53 (2021) 2111–2118. 10.1249/MSS.0000000000002692.

[8] J.R. Fletcher, B.R. MacIntosh, Running Economy from a Muscle Energetics Perspective, Front Physiol 8 (2017) 433. 10.3389/fphys.2017.00433.

[9] B. Van Hooren, P. Teratsias, E.F. Hodson-Tole, Ultrasound imaging to assess skeletal muscle architecture during movements: a systematic review of methods, reliability, and challenges, Journal of Applied Physiology 128 (2020) 978–999. 10.1152/japplphysiol.00835.2019.

[10] P. Ritsche, M.V. Franchi, O. Faude, T. Finni, O. Seynnes, N.J. Cronin, Fully Automated Analysis of Muscle Architecture from B-Mode Ultrasound Images with DL_Track_US, Ultrasound in Medicine & Biology 50 (2024) 258–267. 10.1016/j.ultrasmedbio.2023.10.011.

[11] J.G. Gillett, R.S. Barrett, G.A. Lichtwark, Reliability and accuracy of an automated tracking algorithm to measure controlled passive and active muscle fascicle length changes from ultrasound, Comput Methods Biomech Biomed Engin 16 (2013) 678–687. 10.1080/10255842.2011.633516.

[12] D.J. Farris, G.A. Lichtwark, UltraTrack: Software for semi-automated tracking of muscle fascicles in sequences of B-mode ultrasound images, Comput Methods Programs Biomed 128 (2016) 111–118. 10.1016/j.cmpb.2016.02.016.

[13] J.F. Drazan, T.J. Hullfish, J.R. Baxter, An automatic fascicle tracking algorithm quantifying gastrocnemius architecture during maximal effort contractions, PeerJ 7 (2019) e7120. 10.7717/peerj.7120.

[14] J. Day, L.R. Bent, I. Birznieks, V.G. Macefield, A.G. Cresswell, Muscle spindles in human tibialis anterior encode muscle fascicle length changes, J Neurophysiol 117 (2017) 1489–1498. 10.1152/jn.00374.2016.

[15] T.J. van der Zee, A.D. Kuo, TimTrack: A drift-free algorithm for estimating geometric muscle features from ultrasound images, PLOS ONE 17 (2022) e0265752. 10.1371/journal.pone.0265752.

[16] R. Marzilger, K. Legerlotz, C. Panteli, S. Bohm, A. Arampatzis, Reliability of a semiautomated algorithm for the vastus lateralis muscle architecture measurement based on ultrasound images, Eur. J. Appl. Physiol. 118 (2018) 291–301. 10.1007/s00421-017-3769-8.

[17] O.R. Seynnes, N.J. Cronin, Simple Muscle Architecture Analysis (SMA): An ImageJ macro tool to automate measurements in B-mode ultrasound scans, PLOS ONE 15 (2020) e0229034. 10.1371/journal.pone.0229034.

[18] M. Rana, G. Hamarneh, J.M. Wakeling, Automated tracking of muscle fascicle orientation in B-mode ultrasound images, J Biomech 42 (2009) 2068–2073. 10.1016/j.jbiomech.2009.06.003.

[19] Y. Zhou, Y.-P. Zheng, Estimation of Muscle Fiber Orientation in Ultrasound Images Using Revoting Hough Transform (RVHT), Ultrasound in Medicine and Biology 34 (2008) 1474–1481. 10.1016/j.ultrasmedbio.2008.02.009.

[20] Y. Zhou, J.-Z. Li, G. Zhou, Y.-P. Zheng, Dynamic measurement of pennation angle of gastrocnemius muscles during contractions based on ultrasound imaging, BioMed Eng OnLine 11 (2012) 63. 10.1186/1475-925X-11-63.

[21] G.-Q. Zhou, P. Chan, Y.-P. Zheng, Automatic measurement of pennation angle and fascicle length of gastrocnemius muscles using real-time ultrasound imaging, Ultrasonics 57 (2015) 72–83. 10.1016/j.ultras.2014.10.020.

[22] D.S. Ryan, N. Stutzig, T. Siebert, J.M. Wakeling, Passive and dynamic muscle architecture during transverse loading for gastrocnemius medialis in man, J Biomech 86 (2019) 160–166. 10.1016/j.jbiomech.2019.01.054.

[23] L.G. Rosa, J.S. Zia, O.T. Inan, G.S. Sawicki, Machine learning to extract muscle fascicle length changes from dynamic ultrasound images in real-time, PLOS ONE 16 (2021) e0246611. 10.1371/journal.pone.0246611.

[24] R.E. Kalman, A New Approach to Linear Filtering and Prediction Problems, Journal of Basic Engineering 82 (1960) 35–45. 10.1115/1.3662552.

[25] H.-J. Lee, S. Jung, Gyro sensor drift compensation by Kalman filter to control a mobile inverted pendulum robot system, in: 2009 IEEE International Conference on Industrial Technology, 2009: pp. 1–6. 10.1109/ICIT.2009.4939502.

[26] R.I. Alfian, A. Ma’arif, S. Sunardi, Noise Reduction in the Accelerometer and Gyroscope Sensor with the Kalman Filter Algorithm, Journal of Robotics and Control (JRC) 2 (2021) 180–189. 10.18196/jrc.2375.

[27] B.D. Lucas, T. Kanade, An iterative image registration technique with an application to stereo vision, in: Proceedings of the 7th International Joint Conference on Artificial Intelligence - Volume 2, Morgan Kaufmann Publishers Inc., San Francisco, CA, USA, 1981: pp. 674–679.

[28] R. Duda, P. Hart, Use of the Hough transformation to detect lines and curves in pictures, Commun. ACM 15 (1972).

[29] J. Verheul, S.-H. Yeo, A Hybrid Method for Ultrasound-Based Tracking of Skeletal Muscle Architecture, IEEE Trans Biomed Eng 70 (2023) 1114–1124. 10.1109/TBME.2022.3210724.

[30] J. Shi Tomasi, Good features to track, in: 1994 Proceedings of IEEE Conference on Computer Vision and Pattern Recognition, 1994: pp. 593–600. 10.1109/CVPR.1994.323794.

[31] B.J. Raiteri, L. Lauret, D. Hahn, Residual force depression is not related to positive muscle fascicle work during submaximal voluntary dorsiflexion contractions in humans, J Physiol 602 (2024) 1085–1103. 10.1113/JP285703.

[32] P. Bakenecker, T. Weingarten, D. Hahn, B. Raiteri, Residual force enhancement is affected more by quadriceps muscle length than stretch amplitude, eLife 11 (n.d.) e77553. 10.7554/eLife.77553.

[33] P. Tecchio, B.J. Raiteri, D. Hahn, Eccentric exercise ≠ eccentric contraction, Journal of Applied Physiology 136 (2024) 954–965. 10.1152/japplphysiol.00845.2023.

[34] H.J. Hermens, B. Freriks, C. Disselhorst-Klug, G. Rau, Development of recommendations for SEMG sensors and sensor placement procedures, J Electromyogr Kinesiol 10 (2000) 361–374. 10.1016/s1050-6411(00)00027-4.

[35] A.L. Hof, Jw. Van den Berg, EMG to force processing I: An electrical analogue of the hill muscle model, Journal of Biomechanics 14 (1981) 747–758. 10.1016/0021-9290(81)90031-2.

[36] T. Finni, N.J. Cronin, D. Mayfield, G.A. Lichtwark, A.G. Cresswell, Effects of muscle activation on shear between human soleus and gastrocnemius muscles, Scandinavian Journal of Medicine & Science in Sports 27 (2017) 26–34. 10.1111/sms.12615.

[37] A. Falisse, G. Serrancolí, C.L. Dembia, J. Gillis, I. Jonkers, F. De Groote, Rapid predictive simulations with complex musculoskeletal models suggest that diverse healthy and pathological human gaits can emerge from similar control strategies, J R Soc Interface 16 (2019) 20190402. 10.1098/rsif.2019.0402.

[38] T. Delabastita, M. Afschrift, B. Vanwanseele, F. De Groote, Ultrasound-Based Optimal Parameter Estimation Improves Assessment of Calf Muscle–Tendon Interaction During Walking, Ann Biomed Eng 48 (2020) 722–733. 10.1007/s10439-019-02395-x.

[39] T.J.M. Dick, A.A. Biewener, J.M. Wakeling, Comparison of human gastrocnemius forces predicted by Hill-type muscle models and estimated from ultrasound images, Journal of Experimental Biology 220 (2017) 1643–1653. 10.1242/jeb.154807.

